# Crossvalidation in Brain Imaging Analysis

**DOI:** 10.1101/017418

**Authors:** Nikolaus Kriegeskorte

## Abstract

Crossvalidation is a method for estimating predictive performance and adjudicating between multiple models. On each of k folds of the process, k-1 of k independent subsets of the data (training set) are used to fit the parameters of each model and the left-out subset (test set) is used to estimate predictive performance. The method is statistically efficient, because training data are reused for testing and performance estimates combined across folds. The method requires no assumptions, provides nearly unbiased (slightly conservative) estimates of predictive performance, and is generally applicable because it amounts to a direct empirical test of each model.

## Introduction

Empirical science often involves an informal process of hypothesis generation followed by a formal experimental test of a specific hypothesis. Both hypothesis generation and testing are empirical processes, but the first is exploratory and inconclusive, and the second is confirmatory and supported by formal statistical inference. This two-stage procedure is mirrored in data analysis, where exploratory analyses can help generate specific hypotheses and the latter are subjected to formal inference. Similarly, the fitting of the parameters of a model can convert a vague and difficult-to-test hypothesis into a specific and testable hypothesis.

Crossvalidation is a statistically efficient method for using the cycle of fitting (i.e. generation of a specific hypothesis) and testing within the analysis of a single data set. More specifically, crossvalidation can be used to estimate the predictive performance of alternative models that require parameter fitting. The method can also be used to adjudicate between different models. One element of crossvalidation is the division of the data set into independent subsets used for generation of a specific hypothesis (also known as “parameter estimation”, “fitting” or “training”) and validation (also known as “testing”)^1^. The other element is the use of each data subset for training and testing on different “folds” of the process. This reuse of the data for the purposes of training and testing is what the prefix “cross” in crossvalidation refers to. It renders the process statistically efficient.

Let us first consider the case of two-fold crossvalidation. The data is divided into two subsets, typically of equal size. One set is designated as the training set, the other as the test set. The training could consist in fitting the weights of a linear model. The testing might involve measuring the accuracy of the predictions of the model (e.g. the classification accuracy or the coefficient of determination). More generally, model fitting can be thought of as the generation of a specific hypothesis (the fitted model) from a general space of hypotheses (the model’s parameter space). After using one set of data to generate the hypothesis, using the same set to test the hypothesis would be circular (Kriegeskorte et al., 2009): The specific hypothesis (the fitted model) will reflect the noise to some extent. New data with independent noise can serve to test the specific hypothesis. The use of an independent data set for testing ensures that overfitting does not positively bias the estimate of predictive performance. Using independent data sets simply enables us to perform *an empirical test* of predictive performance.

Using one half of the data for training and the other half for testing provides a valid test. However, we might have designated the training set as the test set and vice versa. Swapping the two sets will give slightly different, but equally valid results. This motivates using each subset as the test set once and averaging the estimates of predictive performance. Note that (a) the training data are independent of the test data on each fold, (b) the test data are independent between folds, and (c) the training data are independent between folds in two-fold crossvalidation. Nevertheless, the estimates of predictive performance obtained on different folds are not independent, because each fold uses all data. For an intuition on this dependency of the results of different folds, consider the effect of the noise on the estimates of predictive performance. If the noise makes data set 1 appear slightly more consistent with data set 2, then predictive performance is likely to appear slightly greater on both folds of crossvalidation. Conversely, if the noise makes data set 1 appear slightly less consistent with data set 2, then predictive performance is likely to appear slightly smaller on both folds of crossvalidation. If the estimates of predictive performance across folds were independent, we would expect averaging of the estimates across folds to reduce the variance of the overall estimate by a factor equal to the square root of the number of folds. Because the folds are dependent, averaging of the estimates across folds improves the estimate by a smaller, but typically still substantial, factor.

There is an obvious disadvantage to two-fold crossvalidation. Although all the data contribute to the estimate of predictive performance, the training set comprises only half the data on each fold. This typically means that the model is not fitted as well as it would be if all data had been used. Our crossvalidation estimate of predictive performance is therefore negatively biased in the sense that it will tend to underestimate the predictive performance our model would achieve on new data if it were fitted with all our data. This motivates the use of a greater portion of the data as training data on each fold. In *k*-fold crossvalidation (Fig. 1), the data is divided into *k* independent subsets. On each fold, one subset is held out as the test set, the others are used for training. As before, the estimates of predictive performance are averaged across folds.

**Figure 1:**
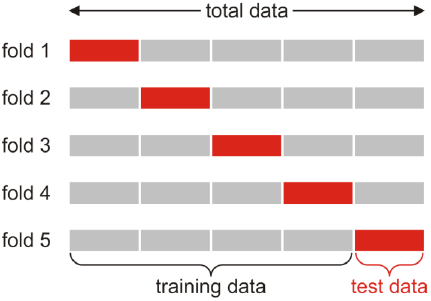
The division of the data into independent training and test sets in 5-fold crossvalidation.

## What’s the optimal number of folds?

Consider the case of *k*=10. In this case, 90% of the data is used for training on each fold. As a result the negative bias of the crossvalidation estimate of predictive performance will typically be small. Although the test set is smaller by factor *k*/2 on each fold (compared to two-fold crossvalidation), there are also *k*/2 times as many folds, across which the evidence will be combined to reduce the variance of the estimate of predictive performance. Using *k*>2, thus, appears statistically advantageous. However, more computation is required, because the model needs to be fitted *k* times.

Beyond the higher computational demands, there is another disadvantage, however, to increasing *k*. For *k*=2, the two training sets are independent between folds. For *k*>2, the training sets are overlapping, and thus dependent, between folds. As explained above, the predictive-performance estimates will always be dependent across folds, even for nonoverlapping training sets. However, their dependence grows with the dependence of the training sets across folds. This reduces the benefit of averaging the performance estimates across folds.

The extreme case is *n*-fold crossvalidation, where *n* is the number of data points. This is also known as leave-one-out crossvalidation. One data point (or the smallest quantum of data on which predictive performance can be tested in a given scenario, e.g. one subject) is held out on every fold. The training sets, then, are identical for all but one data point and the fitted models will be very similar across folds. The estimate of predictive performance will be optimal in terms of its bias (the conservative bias will be minimized), but not in terms of its variance, which will be larger than for smaller settings of *k*. Empirical studies suggest that choices of *k* in the range of 5 to 10, yield good results in practice (Hastie & Tibshirani 2002).

## Important things to keep in mind when using crossvalidation

### (1) Training and test data must be independent

It is essential to ensure that the subsets the data are divided into are statistically independent. For example, fMRI time series are serially autocorrelated in terms of both the underlying signals and the noise. Two time points in the same temporal neighbourhood therefore must never straddle a division into training and test sets. To avoid this, the data can be divided into scanner runs (or subruns, with an appropriate hemodynamic safety margin), and two disjoint sets of runs designated as the training and test data on each fold.

### (2) The folds yield dependent performance estimates

Any inferential procedures must not assume that the performance estimates obtained on different folds are independent. For example, when response patterns are classified and counts of correct and incorrect classifications summed over folds, a binomial test of the null hypothesis that performance is at chance level is not appropriate. A good approach is to obtain a single unbiased performance estimate for each subject, then use a non-parametric test (e.g. Wilcoxon’s signed-rank test) to test whether performance exceeds chance level. This approach can also be extended to model comparisons (using performance differences as the test statistic). It has the additional advantage of treating the variation across subjects as a random effect, and thus supports inferences about the population.

### (3) Multiple nested levels of crossvalidation might be needed

Crossvalidation serves the purpose to obtain estimates of predictive performance that are not biased by overfitting. In many scenarios, there is more than one level of parameters to be fitted. In a linear pattern classification analysis, for example, the classifier might have hyperparameters in addition to the weights associated with each response channel (e.g. each voxel). If the hyperparameters are to be chosen so as to maximise the crossvalidation estimate of performance, the maximum performance estimate will be biased by overfitting of the hyperparameters. A second level of crossvalidation is therefore needed. On each fold of the outer crossvalidation loop, the hyperparameter can be determined by nested crossvalidation within the training set. The optimal setting is then used to predict the test set (which has not been used to optimise the hyperparameter).

## Examples of brain imaging analyses that do or do not require crossvalidation

### Example 1: Univariate activation analysis

The effects of experimental conditions on activation within a predefined brain region are often analysed with linear models. The framework of univariate linear regression does not require us to divide the data or perform crossvalidation. In the analysis of functional magnetic resonance imaging data, for example, it is typically reasonable to assume that the errors are univariate normal and to account for temporal autocorrelation of the time series by pre-whitening or pre-colouring. The predictor weights and any linear contrasts of interest, along with their standard errors, can then be estimated and inference can be performed using a single data set. This approach is generally preferred to a crossvalidation approach for its power, elegance, and convenience.

Even in univariate activation analysis, however, the use of independent validation sets can be essential. For example, when a given data set is used to explore the brain for locations exhibiting a particular effect (to define regions of interest) and statistically related activation effects are then to be analysed for the region thus defined. In this scenario, independent data are needed to ensure valid statistical inference and unbiased estimates of effect sizes (Kriegeskorte et al. 2009; Vul et al. 2009; Kriegeskorte et al. 2010). Independent validation data are also needed when a large number of univariate activation predictors is fitted with regularisation, and when the model’s ability to generalize to a different set of stimuli is to be tested (e.g. Kay et al. 2008; Huth et al. 2012).

### Example 2: Univariate activation mapping

In the previous example, we assumed a single predefined region of interest on whose regional-average activation was analysed by univariate regression. Let us consider the brain mapping problem. The hypothesis that contrasting two cognitive states will reveal greater activity *somewhere* in the brain is more difficult to test than the hypothesis that there will be brain activity in a specific region. We need to analyse all locations and perform inference on the map as a whole. The field has developed powerful techniques for mapping activation contrasts throughout a measured volume, while accounting for temporal and spatial dependencies, and for the multiple testing across locations. One approach relies on Gaussian field theory (Friston et al. 1994; Worsley et al. 1996), another uses nonparametric permutation tests (Nichols & Holmes, 2002; see also article 324. Non-parametric procedures). Both of these aim to control the familywise error rate (Nichols & Hayasaka 2003). An alternative approach to the multiple-testing problem is to control the false-discovery rate (Genovese et al. 2002; see also article 323. False discovery rate procedures). None of these approaches require crossvalidation. Classical brain mapping is an example of a scenario where the complication of crossvalidation can be avoided. Many of the established methods rely on particular assumptions (e.g. univariate normal errors), which are justified and which increase the power of the statistical inference.

### Example 3: Multivariate pattern-information analysis

Analyses of multivariate activity patterns within a predefined region of interest can reveal the information contained in local brain representations (for a tutorial introduction, see Mur et al. 2009). One approach to detecting information in patterns would be multivariate linear regression (Krzanowski 1988), the multivariate generalization of the analysis framework used for univariate activation analysis. However, this framework relies on the assumption that the errors are normally distributed. Whereas the normal assumption is reasonable for the univariate scenario, it is not in general safe to rely on multivariate normal errors in pattern analyses (Kriegeskorte at al., 2006). This is one reason why the field has preferred permutation tests and crossvalidation-based approaches including linear discriminant analysis. When a Fisher linear discriminant or linear support-vector machine (Bishop 2006) is fitted with one data set and its performance compared to chance level using independent data, the assumptions of the model are implicitly tested. For example, the Fisher linear discriminant is based on a multivariate normal model of the errors, much like the multivariate regression framework. However, a violation of multinormality would only decrease predictive performance. Inference on crossvalidated performance estimates therefore provides a valid test of pattern information that does not rely on the multinormal assumption for its validity (although it does depend on multinormality for optimal power). This illustrates how crossvalidation can reduce our reliance on questionable assumptions.

### Example 4: Multivariate pattern-information mapping

As for univariate activation, it is useful to map an imaged volume continuously so as to locate brain regions containing a particular kind of information in their local multivariate patterns (Kriegeskorte et al. 2006). One approach relies on nonparametric permutation tests (as used in univariate mapping), where the stimulus labels are permuted to simulate the null hypothesis. Such tests do not require distributional assumptions and enable us to use the most suitable test statistics including those provided by classical multivariate statistics (Krzanowski 1988), measures of the similarity structure of the patterns (Kriegeskorte et al. 2008), or crossvalidated pattern-classifier performance estimates. Another approach is to enter multiple single-subject descriptive pattern-information maps into a multisubject inferential mapping procedure, treating subject as a random effect. As for the permutation approach, the pattern-information statistics may or may not involve crossvalidation.

### Example 5: Selection among multiple nonlinear computational models

A major goal of brain science is to test computational models of brain information processing. Brain imaging studies have begun to incorporate such models into the data analysis (e.g. Kay et al. 2008; for a review see Kriegeskorte 2011). In contrast to the generic statistical models discussed in the previous examples, a model of this type mimics the actual information processing performed by the brain. For example, it may take stimuli in the form of bitmap images as its input and predict their representation in a visual area. Interesting models of brain information processing are typically complex and nonlinear. In order to test and compare models, we need to determine the extent to which they can predict brain representations of arbitrary novel stimuli (or novel stimuli within a predefined population of stimuli). When models of this type have parameters to be fitted, the use of independent validation sets (data for different stimuli) is essential. Current analyses of this type typically require crossvalidation procedures.

## What are the alternatives to crossvalidation and what are their advantages and disadvantages?

### Bayesian model selection

Evaluation of the predictive performance of a model and selection among alternative models can be approached using probability theory. The predictive performance of a model m on a data set d can be measured as p(d|m), the probability of the data given the model. The philosophically most compelling way to select a model is by its probability given the data, p(m|d), which can be computed using Bayes theorem (MacKay 2003; Bishop 2006; see also articles 325. Bayesian model inversion and 328. Bayesian model selection). A fully probabilistic treatment of model selection requires explicit representation of all sources of uncertainty including a prior probability distribution over the model space. This approach requires no fitting. It therefore does not suffer from overfitting, and does not require empirical validation on independent data (Ghahramani 2013). A fully probabilistic approach, while simple in theory, can be daunting in practice. First, modelling all sources of uncertainty is often difficult, and results will depend on the assumptions made. A second challenge is probabilistic inference. A growing literature on stochastic and deterministic approximate inference algorithms, including Markov chain Monte Carlo sampling, provides powerful and general solutions to this challenge. However, the particular problem at hand may require a custom-built solution for inference. Fully automatic methods are not yet widely accessible. Fitting a model’s parameters greatly simplifies inference and can render an otherwise untestable model testable. Therefore empirical tests on independent data (as efficiently implemented by crossvalidation) remain an important tool for scientific data analysis – whether the inference framework adopted is Bayesian or frequentist, or a combination of both. Assumptions implicit to the models, then, are continually confronted with the ultimate challenge: predicting new data.

### Minimum description length

Another criterion for model selection is the minimum description length (MDL) principle. The MDL principle states that the best model is the one that enables the most efficient compression of the data. The compressed data must describe the model, its parameters, and the residuals of the fit to the data. Complex models are implicitly penalized because their description takes more space. The winning model best balances its own complexity against the size of the residuals (and thus the number of bits needed to transmit them at a specified precision). The preference for models that can be concisely described can be interpreted as a prior favouring simple models. The MDL principle then turns out to be equivalent to Bayesian inference (MacKay 2003). In practice, determining the MDL by optimally encoding models, parameters, and residuals for compressed transmission is a nontrivial engineering challenge. The explicitly Bayesian approach may be easier to implement and preferable for its conceptual clarity.

### Information criteria

In certain scenarios, we can avoid both the challenge of a fully Bayesian approach and the computational demands of crossvalidation. The Akaike information criterion (AIC) and the Bayesian information criterion (BIC) provide measures of model performance that account for model complexity. AIC and BIC combine a term reflecting how well the model fits the data with a term that penalizes the model in proportion to its number of parameters. These criteria are easier to compute than a crossvalidation estimate of predictive performance and they enable accurate model selection when the assumptions they are based on hold. The AIC relies on an asymptotic approximation that may not hold for a given finite data set, and the BIC relies on the assumption that the model errors are independent and normally distributed.

Both AIC and BIC are functions of the parameter count and the maximised likelihood, i.e. the probability of the data given the maximum-likelihood fit of the model. Counting parameters is not in general a good method of estimating model complexity. For example, the effective number of parameters is reduced when the hypothesis space is regularised using an explicit prior or by including a penalty on undesirable parameter combinations in the cost function minimised by the fitting procedure. The effective number of parameters can be difficult to estimate accurately. Nevertheless, where applicable, AIC and BIC provide a quick and easy way to compare models.

## Concluding remarks

Statistical inference, whether Bayesian or frequentist, necessarily combines data with (explicit or implicit) prior assumptions. Prior assumptions can stabilise our estimates and guide our inferences. In Bayesian inference, a prior – when correct – can improve the accuracy of our inference. In frequentist inference, similarly, prior assumptions (e.g. about the error distribution) – when correct – can lend us power. Unsurprisingly, inference techniques that make fewer assumptions tend to have less power.

The classical statistical approach is to decide on a set of prior assumptions and then fit and perform inference on the basis of a single data set. In theory, this obviates the need for checking predictive performance on independent data. In a fully Bayesian framework, overfitting is not an issue. In a frequentist framework, overfitting can be accounted for analytically for simple models (e.g. when we compute the standard error of a parameter estimate for a linear model with independent normal errors). A likelihood-ratio test comparing two point hypotheses has been shown to be optimally powerful (Neyman & Pearson 1933) – and thus more powerful than crossvalidation-based tests – when its assumptions hold. This reflects the benefits of prior knowledge.

Without tests of predictive performance on independent data, however, the classical statistical approach to inference is severely limited, for two reasons. First, our assumptions are usually not exactly true, and therefore our inferences are not necessarily reliable. Second, the classical statistical approach is only feasible for a very restricted class of models. This is the reason why the field that has led the development of the most complex models, machine learning, heavily relies on crossvalidation.

In science our models should mirror the mechanisms we hypothesise, and not be limited to a small set we happen to know how to test with a single data set. Our goal is not mathematical elegance, but learning about nature. Testing effects and selecting models according to their actual predictive power on new data puts all assumptions to the test and keeps us firmly grounded in empirical reality.

In sum, the advantage of crossvalidation over alternative methods is its generality: It can be applied when other methods cannot and it does not rely on assumptions or approximations. For many of the most interesting and well-motivated models in brain science, a fully Bayesian approach is daunting and the assumptions required for classical frequentist inference and for information criteria for model selection may not hold. Crossvalidation enables us to develop our models as motivated by the science (rather than the statistics) and to employ the familiar procedure of first defining a hypothesis specific enough to be testable and then testing it empirically within the analysis of a single data set.

## GLOSSARY

Generalization performance: the quality of the predictions about new data afforded by a model fitted with a given data set.
Overfitting: the inevitable effect of measurement error on the estimates of parameters obtained by fitting a model to a given data set.
Independence (statistical independence): the absence of any relationship, linear or nonlinear, deterministic or stochastic, between two variables. Independence implies that learning either variable does not change our belief (expressed as a probability distribution) about the other variable.

## Footnotes

1 Some authors use the term *validation set*, and reserve the term *test set* for a third independent subset of the data, used when a second level of validation is required. Here we use the concepts of validation and test interchangeably.

